# GraphGR: A graph neural network to predict the effect of pharmacotherapy on the cancer cell growth

**DOI:** 10.1101/2020.05.20.107458

**Authors:** Manali Singha, Limeng Pu, Abd-El-Monsif Shawky, Konstantin Busch, Hsiao-Chun Wu, J. Ramanujam, Michal Brylinski

**Affiliations:** Department of Biological Sciences, Louisiana State University, Baton Rouge, LA 70803, USA; Center for Computation and Technology, Louisiana State University, Baton Rouge, LA 70803, USA; Department of Cell Biology, National Research Centre, 12622 Giza, Egypt; Division of Computer Science and Engineering, Louisiana State University, Baton Rouge, LA, 70803, USA; Division of Electrical and Computer Engineering, Louisiana State University, Baton Rouge, LA, 70803, USA

**Author notes:** Contributed equally (Manali Singha, Limeng Pu).

**Keywords:** cancer growth rate, kinase inhibitors, differential gene expression, gene-disease association, cancer-specific networks, network biology, graph neural network, graph reduction, propagation attention, artificial intelligence

## Abstract

Genomic profiles of cancer cells provide valuable information on genetic alterations in cancer. Several recent studies employed these data to predict the response of cancer cell lines to treatment with drugs. Nonetheless, due to the multifactorial phenotypes and intricate mechanisms of cancer, the accurate prediction of the effect of pharmacotherapy on a specific cell line based on the genetic information alone is problematic. High prediction accuracies reported in the literature likely result from significant overlaps among training, validation, and testing sets, making many predictors inapplicable to new data. To address these issues, we developed GraphGR, a graph neural network with sophisticated attention propagation mechanisms to predict the therapeutic effects of kinase inhibitors across various tumors. Emphasizing on the system-level complexity of cancer, GraphGR integrates multiple heterogeneous data, such as biological networks, genomics, inhibitor profiling, and genedisease associations, into a unified graph structure. In order to construct diverse and information-rich cancer-specific networks, we devised a novel graph reduction protocol based on not only the topological information, but also the biological knowledge. The performance of GraphGR, properly cross-validated at the tissue level, is 0.83 in terms of the area under the receiver operating characteristics, which is notably higher than those measured for other approaches on the same data. Finally, several new predictions are validated against the biomedical literature demonstrating that GraphGR generalizes well to unseen data, i.e. it can predict therapeutic effects across a variety of cancer cell lines and inhibitors. GraphGR is freely available to the academic community at https://github.com/pulimeng/GraphGR.

## Introduction

Cancer initiation and progression involve a sequence of gene-environment interaction events changing the gene expression and ultimately leading to the disruption of homeostasis [1]. The phosphorylation of various proteins is one of the key processes regulating various cellular functions, including cell cycle, apoptosis, proliferation, differentiation, growth, and others. The phosphorylation of tyrosine, serine, and threonine residues is the primary function of kinase proteins [2], 538 of which are encoded by the human genome [3]. Any disruption of kinase activity can trigger the dysregulation of cellular functions and many dysregulated kinases have oncogenic effects responsible for cancer progression [2]. The discovery of kinase inhibitors for cancer therapy has changed the course of treatment from a conventional chemotherapy to the targeted pharmacotherapy. Although selective inhibitors are available to target certain kinases in human cancers [4], the majority of compounds bind to the highly conserved ATP binding sites of multiple targets [5–7]. Certainly, the binding promiscuity of kinase inhibitors can lead to adverse drug reactions [3, 8, 9], but also to the desired polypharmacological effects by simultaneously targeting multiple proteins involved in cancer-related processes [10–12]. Large-scale kinase inhibitor profiling data providing the information on the inhibition of enzymatic activity across the human kinome [13, 14] greatly facilitate research on kinase-centric polypharmacological anticancer agents [15–17].

Numerous studies demonstrate that the accumulation of genetic alterations and the subsequent changes in gene expression patterns are major factors driving cancer progression [18–20]. The identification of differentially expressed genes not only enhances our understanding of cancer biology [21–23], but it also reveals new targets for therapeutic intervention [24–26]. Many methods combine genomics with drug chemical and activity information to predict the response to drugs in cancer treatment. For instance, CDRscan predicts anticancer drug responsiveness based on drug screening assay data, genomic profiles of human cancer cell lines, and molecular fingerprints of drugs [27]. The analysis of the observed and predicted drug response showed an exceptionally high accuracy of CDRscan with a mean coefficient of determination of 0.84 and the area under the receiver operating characteristics (ROC) of 0.98. Another method, DeepDR, predicts drug response purely based on the mutation and expression profiles of cancer cells. The reported overall prediction performance of DeepDR is also exceptionally high with a mean squared error of only 1.96 in the log-scale IC_50_ values. Further, SRMF predicts anticancer drug responses of cell lines solely from the chemical structures of drugs and the baseline gene expression levels in cell lines [28]. Those two features are used as regularization terms, which are incorporated into the drug response matrix factorization model. SRMF yields a drug-averaged mean squared error of 1.73 between predicted and observed responses of sensitive and resistant cell lines. Nonetheless, the performance of all these algorithms is likely grossly overestimated on account of a random split of the redundant data into training, validation, and testing subsets resulting in significant overlaps among these sets.

The problem at hand is actually a bit more complicated on account of the high intricacy of cancer mechanism. In contrary to the abovementioned studies, the drug responsiveness is unlikely to be accurately predicted from drug molecular structures, generic binding assays, and genetic features of cancer cells using standard convolutional neural networks, which have been developed primarily for applications within the Euclidean domain. Carcinogenesis is, in fact, a systems-level, network phenomenon with a complex phenotype heavily depending on the flow of signaling information within a cell [29–31]. Biological networks, including protein-protein interaction (PPI) networks, are often employed to study alterations in information flow in cancer cells caused by oncogenic changes in protein activity and expression [32–34]. For instance, methods utilizing the information flow in networks can extract the functional diseasegene relationships in order to prioritize candidate genes for pharmacotherapy [35]. Therefore, network-based techniques are generally more suitable to examine the overall effects of drugs on the cancer cell growth. Nonetheless, these approaches require geometric deep learning systems applicable to domains such as graphs, point clouds, and manifolds [36].

Many non-Euclidean, real-life data such as social networks, communication networks, biological networks, molecular structures, etc., can be represented as graphs. Yet, very little attention has been devoted to study the graph information processing in terms of machine learning systems until the last few years. One of the earliest graph neural networks (GNNs) utilizes the graph structure to learn the representation of the input data [37]. The major limitation of this method is that it restricts the information propagation to the first-order neighbors of every node limiting the information flow in the model. More recently, a graph convolutional network (GCN) was proposed to provide a more flexible model propagating information through many orders of neighbors [38, 39]. More advanced models were developed following the fundamental work on GCN, including a graph-based neural network employing the long-short term memory (LSTM) to carry out the information propagation that was demonstrated to have a significantly improved performance [40]. Another information propagation scheme aggregates the average embeddings of the neighboring nodes yielding a high performance especially for node classification in large graphs [41]. Numerous other techniques implementing minor improvements and changes are available to operate on the graph-structured data [42–44].

In this communication, we first describe a procedure to construct information-rich, cancer-specific graphs integrating heterogeneous data on differential gene expression, kinase inhibitor profiling, protein-protein interactions, and disease association scores. Next, we present GraphGR, a GNN-based algorithm employing multiple graph convolutional blocks with attention-based propagation and a sophisticated graph readout mechanism to predict the effect of a drug treatment on the cancer cell growth. The performance of GraphGR is compared to other deep learning, graph kernel, and traditional approaches, in carefully designed crossvalidation benchmarks. Finally, GraphGR is applied to unseen data and selected cases are validated against the biomedical literature.

## Results

### Integration of heterogeneous data to construct cancer-specific networks

Input for GraphGR are cancer-specific networks assembled from multiple heterogeneous data. Data integration was performed by mapping differential gene expression, disease-gene association scores, and kinase inhibitor profiling onto the human PPI network. This procedure is schematically shown in Figure 1. For example, for a certain cell line, up- (green) and down-regulated (red) genes are marked according to the differential gene expression for that cell line and some nodes are also assigned disease-gene association scores (numbers in bold). If this cell line is treated with a kinase inhibitor, pIC_50_ values against its targets are then added to the graph (numbers in italics). In the resulting graph, kinase nodes have gene expression values, and some kinase nodes also have pIC_50_ values and disease-gene association scores, whereas non-kinase nodes have gene expression values and some non-kinases nodes also have diseasegene association scores. Note that all cell line-drug combinations have the same underlying PPI network, however, different cell lines usually have different gene expression values as well as disease association scores. Similarly, different drugs target usually inhibit different sets of kinases, therefore, node features are generally unique for various cell line-drug combinations.

**Figure 1.**
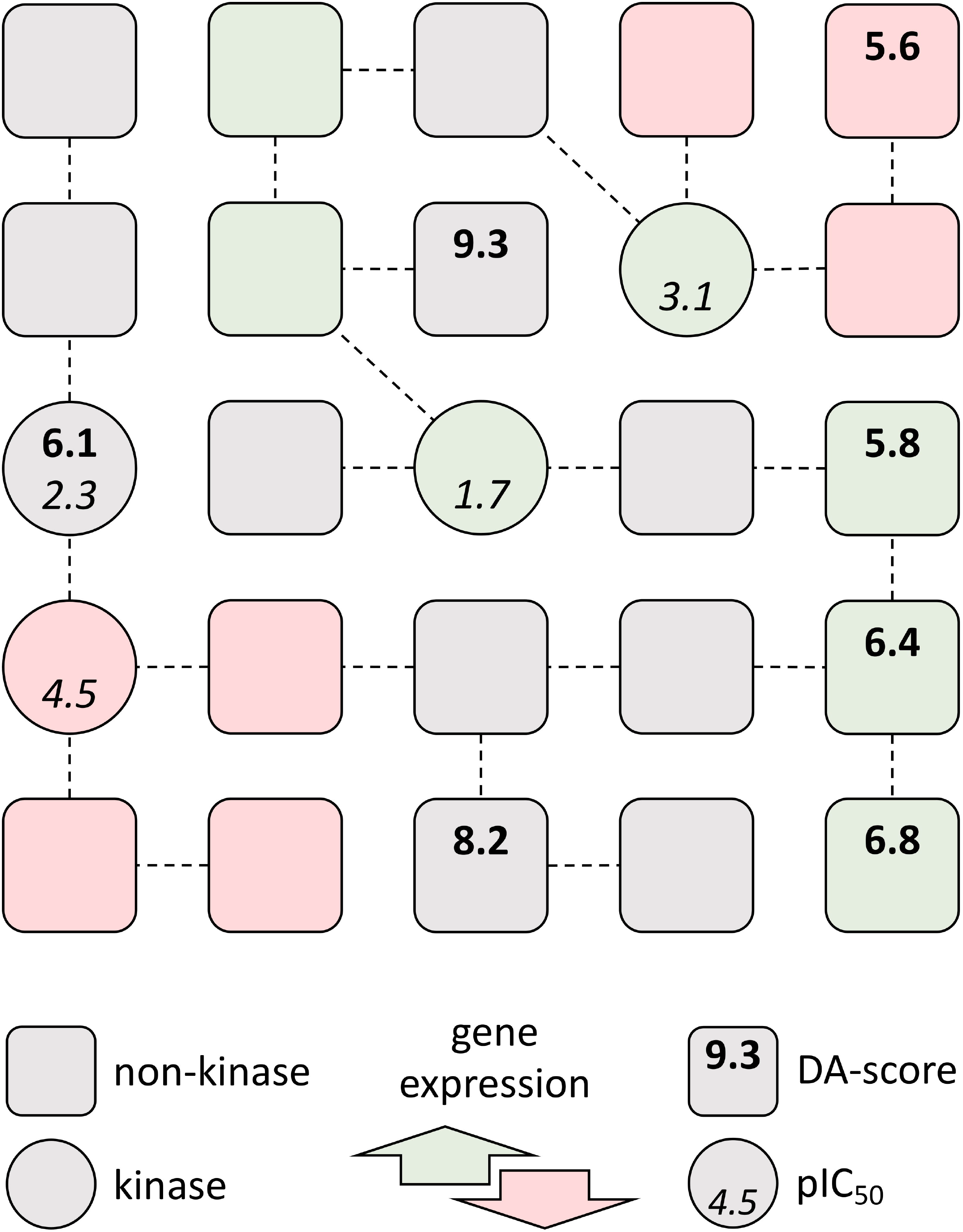
Schematic of the graph representation of multiple heterogeneous data. Circles are kinases, whereas rounded squares represent non-kinase proteins. Nodes are connected through confident interactions forming a graph. Each node is colored according to the differential gene expression: green – up-regulated, red – down-regulated, and gray – normally regulated. Both types of nodes can have gene-disease association scores (numbers in bold), whereas kinases can also have pIC_50_ values according to the kinase profiling data (numbers in italics).

### Analysis of full cancer-specific networks

After initial experiments with full size graphs corresponding to the original PPI networks, we found that there are three issues with this data representation. First, all instances share exactly the same graph topology and differences are only in node features, i.e. gene expression, disease association, and pIC_50_ values, which makes it difficult for the model to gather the information required for effective learning. Second, full size graphs are very sparse with a density (see Eq. 1) as low as 0.004 wasting the computer memory and significantly extending learning time. A quick fix to this problem could be to employ a sparse representation, such as the coordination (COO) format [45], however, this approach is not workable due to the GNN library implementation not supporting sparse data formats. Third, the majority (98%) of nodes in the graph are non-kinase proteins with no inhibition data and most proteins are normally regulated according to the differential gene expression leading to the significant sparsity of important features. As result, most of entries in the feature matrix carry no effective information resulting in a poor learning performance. An illustrative analogy is trying to train a model to classify images having less than 1% different pixels. Given such tiny differences, any model is going to struggle learning the underlying patterns.

### Knowledge-based reduction of cancer-specific networks

In order to address these issues, we devised a knowledge-based graph reduction procedure by edge contraction, which is a fundamental operation in graph theory [46]. Here, the idea is to remove a particular edge and then merge the incident nodes of that edge into a new node. Edge contraction is widely used in recursive formulas for calculating the number of spanning trees of an arbitrary connected graph [47], and in recurrence formulas for calculating the chromatic polynomial of a simple graph [48]. Nonetheless, a simple edge contraction based solely on the connectivity is not going to produce the desired outcome in our case because we also need to account for the features of nodes. The knowledge-based edge contraction employs both connectivity and biological feature information to satisfy the following conditions: both incident nodes are non-kinase proteins, share the same differential gene expression, and belong to the same biological process cluster. The last condition is very important to ensure that the reduction merges only those nodes belonging to the same pathway, thus supporting the biological knowledge. Biological processes in cancer-specific networks are determined by clustering nodes according to the similarity of their Gene Ontology (GO) terms.

GOGO is a method to calculate semantic similarities between GO terms using Directed Acyclic Graphs (DAGs) [49]. GO consists of three DAGs created based on molecular function (MF), cellular component (CC), and biological process (BP) ontologies [50]. In order to verify that the network locality is preserved when using GOGO similarities derived from the biological process ontology, we first calculated similarity values between 1^st^, 2^nd^, 3^rd^, and 4^th^ order neighbors in the full PPI network. Figure 2 shows that GOGO similarities are the highest for the 1^st^ order neighbors and decrease with the increasing order. These results corroborate previous studies demonstrating that the closer the two proteins are in the network the more similar are their biological functions [51]. Next, using GOGO similarities and the hierarchical clustering analysis (HCA), all proteins in the graph were partitioned into 30 (HCA-30), 100 (HCA-100), and 300 (HCA-300) clusters. Only those nodes belonging to the same cluster were allowed to be merged during graph reduction.

**Figure 2.**
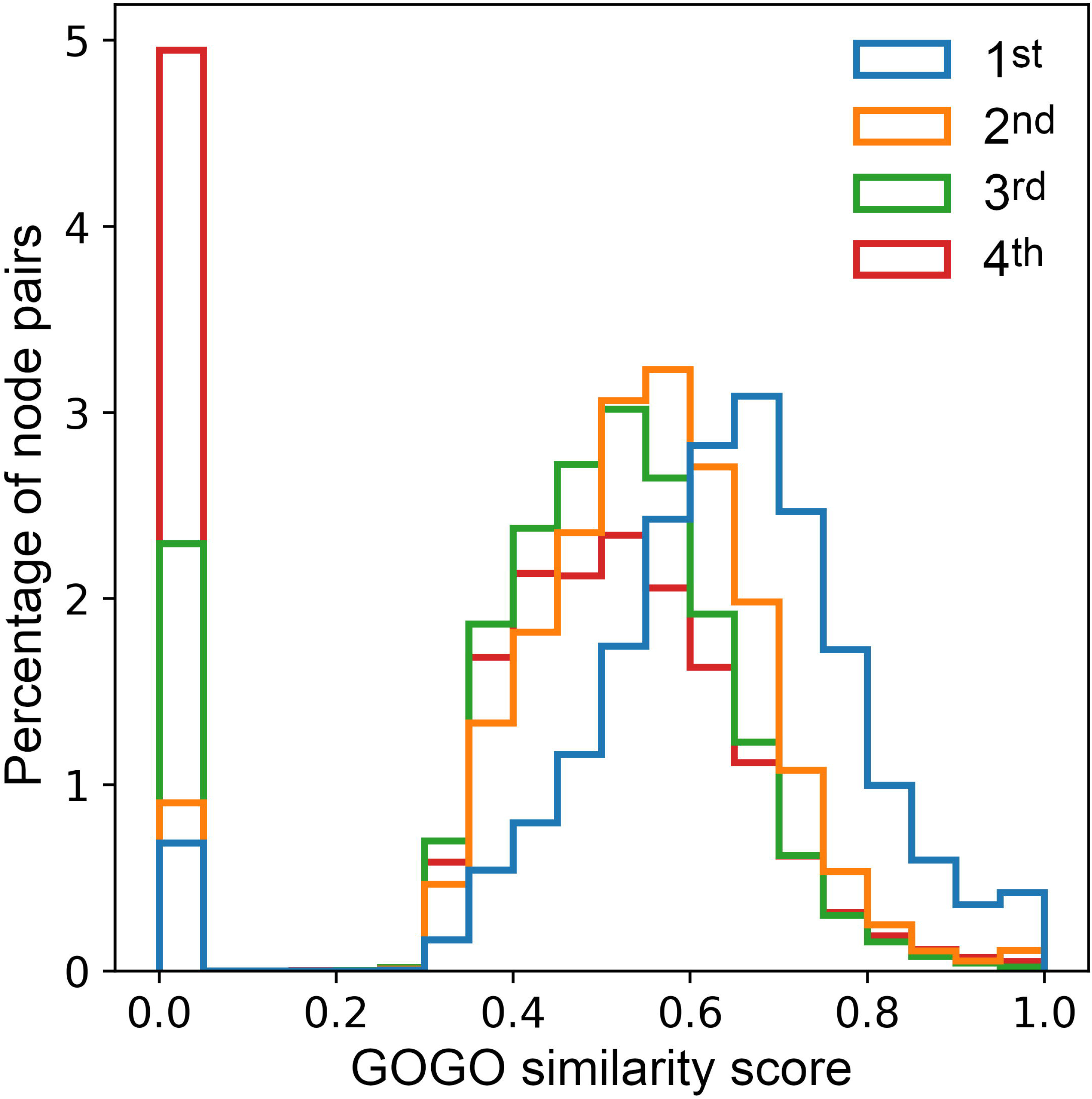
Histogram of the pairwise GOGO similarity scores across the protein-protein interaction network. GOGO similarities are calculated using the biological process ontology for 1^st^, 2^nd^, 3^rd^, and 4^th^ order neighbors in the network.

The graph reduction procedure and example graphs are shown in Figure 3. An edge may be contracted only if both incident nodes are non-kinases, share the same differential gene expression, and belong to the same GOGO cluster. Yellow squares in Figure 3A delineate groups of nodes that can be merged by contracting edges connecting them. The resulting reduced graph shown in Figure 3B has the same number of kinases (circles), but fewer non-kinase proteins (rounded squares and diamonds representing merged nodes). The rationale behind this procedure is not only to reduce the size of a graph, but also to create more diversity among cell lines, which is highly beneficial for further machine learning applications. Note that the reduction is performed on different cell lines without any drug information because we consider only the differential gene expression, the type of node (kinase or non-kinase protein), and the biological process assignment of nodes, with the latter two properties being independent on the cell line type. After the reduction, the average number of nodes decreased from 19,144 to 1,349 and the average graph density (see Eq. 1) increased from 0.004 to 0.014. Compared to the original PPI network (Figure 3C), the reduced graph (Figure 3D) has a much higher ratio of kinase (red dots) to non-kinase (green dots) proteins.

**Figure 3.**
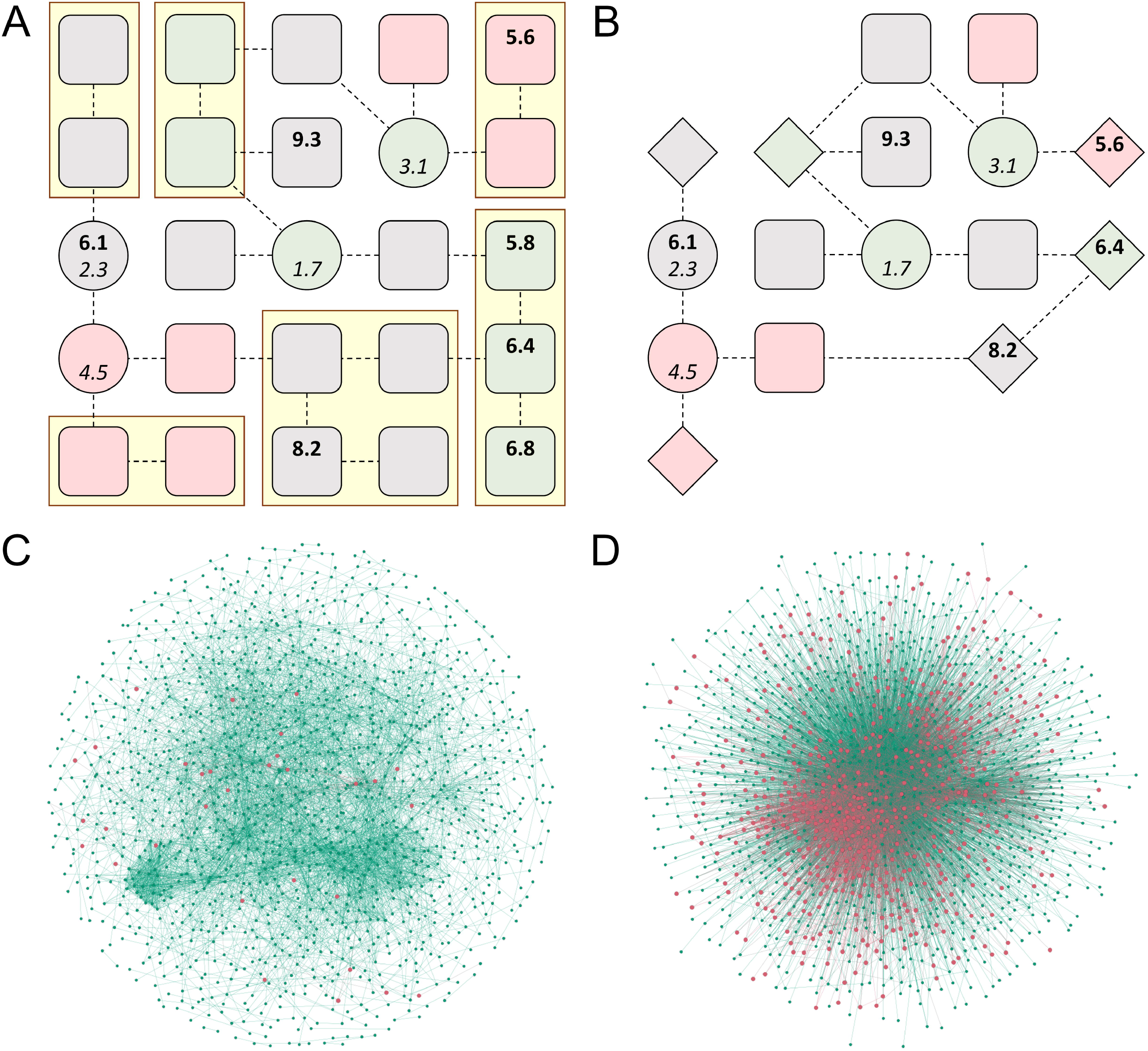
Graph reduction of cancer-specific networks. **(A)** A schematic of the initial graph with yellow boxes outlining groups of nodes that can be merged by contracting their edges. (B) A schematic graph of the reduced graph in which merged nodes are represented by diamonds. **(C)** The initial (sub)network for glioblastoma (cell line A172) with red nodes representing kinases and green nodes representing other proteins. (D) The reduced network for glioblastoma colored the same as in **C.** The network in **C** is a randomly sampled subgraph from the original network with the same number of nodes as D.

### Analysis of reduced cancer-specific networks

The graph reduction procedure greatly increases the diversity of graph topologies and features across all 359 cancer cell lines, while maintaining the important information and biological knowledge in each graph. In order to further demonstrate the effectiveness of the reduction scheme, we calculated information gain/loss after the reduction using the Shannon entropy of features and the graph-feature entropy (see Eq. 3). Ideally, the graph reduction should increase the information content for features without any decrease in graph-feature information. The information gain/loss is shown in Figure 4 for several reduction schemes. Although the simplest reduction requiring incident nodes to share at least one GO-BP term increases the feature-only entropy by 2.3 ±0.6 (green bar), it causes a detrimental decrease of the graph-feature entropy by −0.4 ±0.04 (red bar). In contrast, HCA increases the feature-only entropy while preserving the valuable graph-feature information. In particular, partitioning nodes into 30 clusters not only yields the highest information gain for features of 3.5 ±0.9, but also slightly increases the graph-feature entropy. Based on these results and due to the fact that there are actually 30 level-1 biological processes in GO [52], we incorporated HCA-30 into the graph reduction procedure.

**Figure 4.**
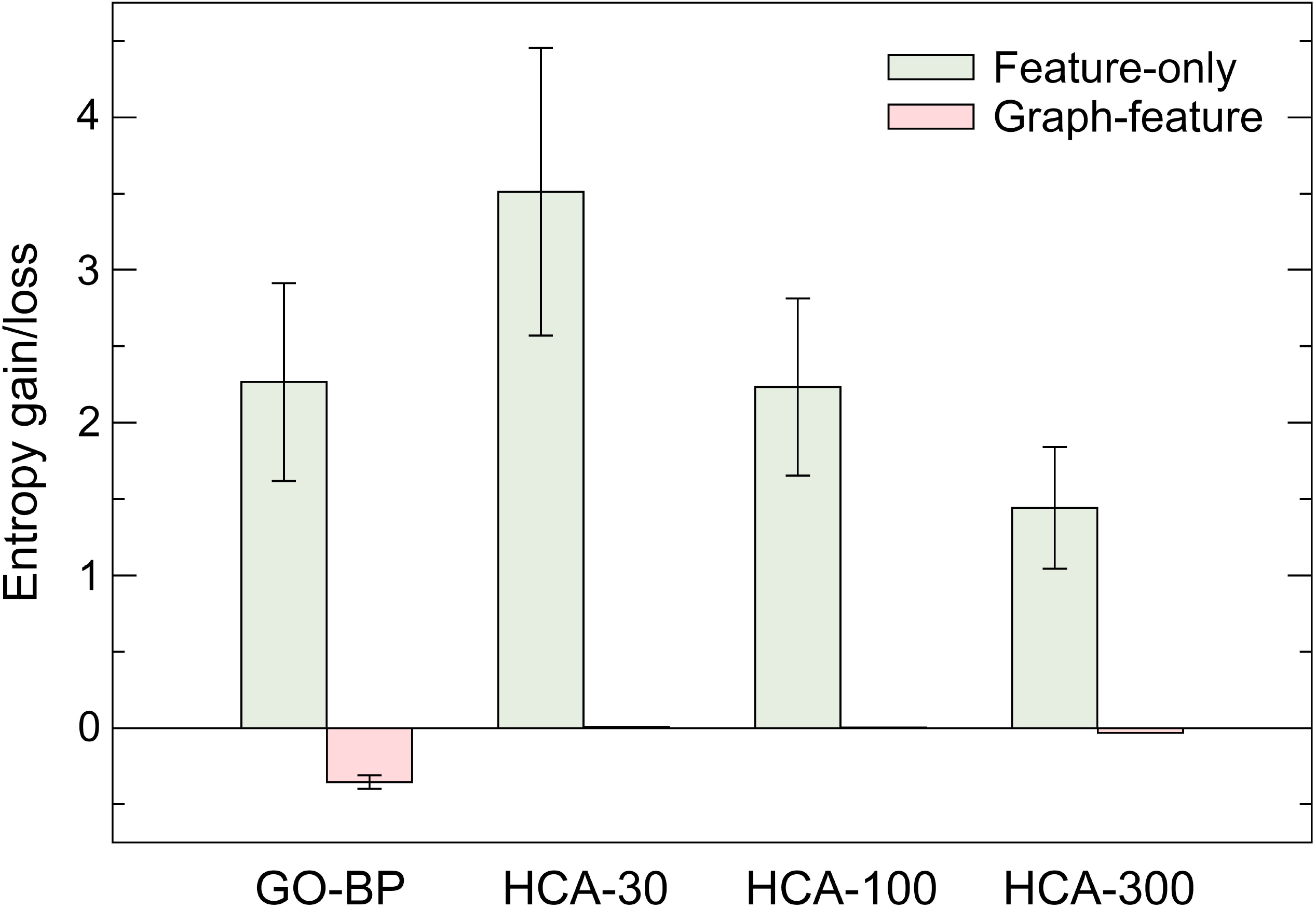
Entropy gain/loss for different reduction schemes. Green bars represent the Shannon entropy calculated using the feature matrix only, while red bars correspond to the graph-feature entropy computed using both feature and topological information of a graph. GO-BP requires that two incident nodes have a common biological process term to be assigned to the same cluster. HCA bars correspond to the clustering using GOGO similarities into 30, 100, and 300 clusters.

### Information propagation in GraphGR

GraphGR implements a GNN model to predict the response of various cancer cell lines to the treatment with different kinase inhibitors. The GNN employs graph convolutions, which are functionally equivalent to matrix convolutions in the CNN working with images. Similar to the CNN propagating the information of a pixel to its neighbor pixels, the GNN propagates the information of a node in the graph to its neighbor nodes. The architecture of GraphGR is presented in Figure 5. An instance consisting of the combination of a cell line and a drug is used to create a cancer-specific network, which is subsequently subjected to the reduction procedure (Figure 5A). The reduced graph is then processed through a cascade of graph convolution blocks (Figure 5B). Each block contains three components, the attention-based propagation, the embedding update, and the generation of new embeddings. Although only the information from 1^st^ order neighbors is passed between nodes in a single block, using multiple sequential blocks actually propagates the information from higher order neighbors.

**Figure 5.**
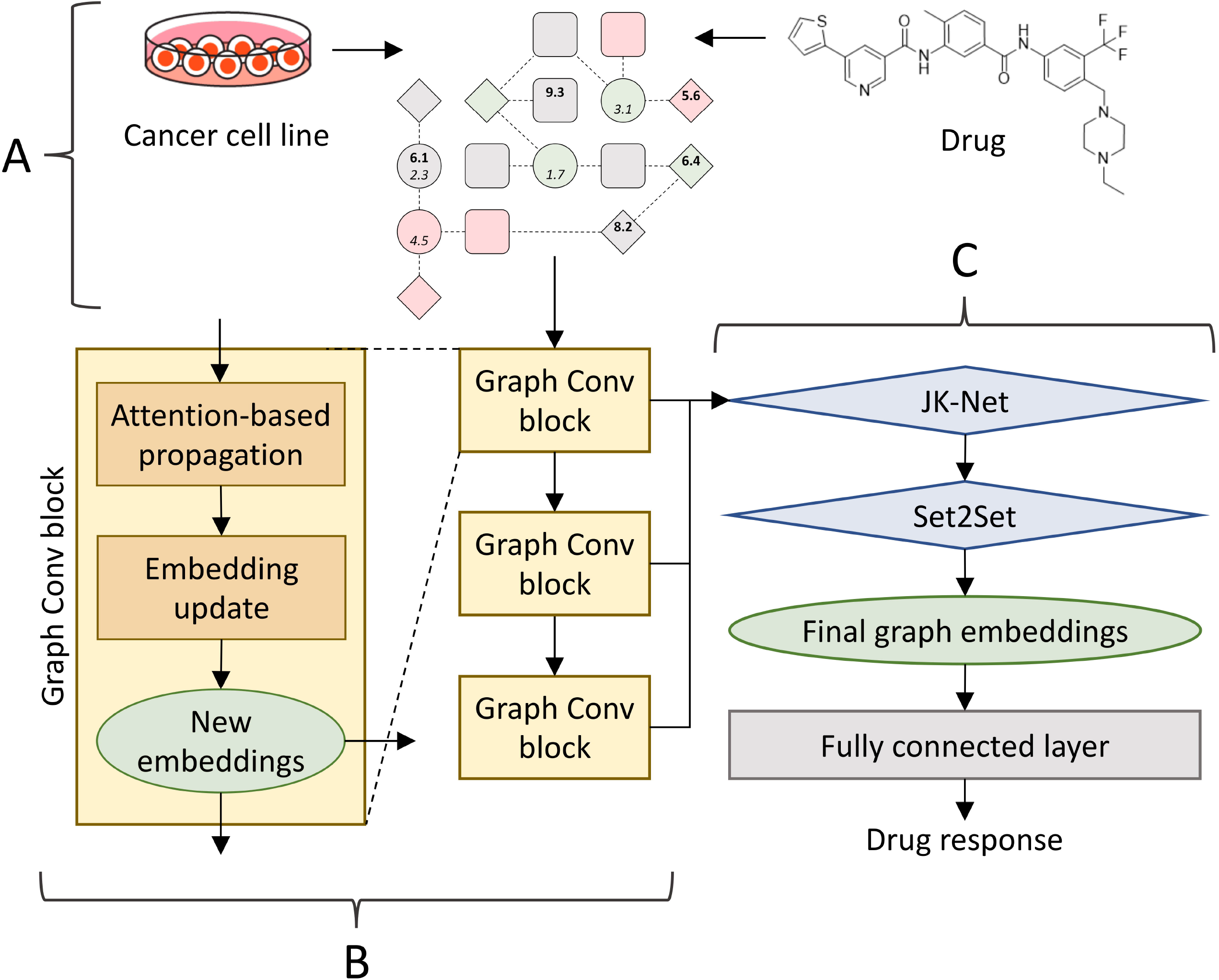
GraphGR architecture. **(A)** The input is a reduced graph constructed for a combination of a cell line and an inhibitor. (B) The graph is processed through a cascade of three graph convolutional blocks. In each block, first an attention-based propagation is utilized to pass the information among nodes, and then a graph isomorphism network is employed to update the embeddings for each node. (C) Node embeddings generated by all blocks in B are combined using a JK-Net layer and passed to a Set2Set pooling layer serving as the read-out function to acquire the final graph embeddings. At the end, graph embeddings are sent to a fully connected layer to predict the drug response.

This procedure is illustrated in Figure 6 for a simple 4-node graph. Initially, each node has its own information (color coded in Figure 6A), which is used to generate node embeddings. In our model, nodes are proteins connected through PPIs and the information comprises gene expression, disease association, and pICso values. During the first propagation step, a node of interest, such as node 1 in Figure 6, receives information from its 1^st^ order neighbor, node 2 (Figure 6B). At the same time, node 2 receives information from its 1^st^ order neighbors, nodes 3 and 4. Node 1 now contains more information that is used to generate new embeddings. In the second propagation step, the information from nodes 3 and 4 already present in node 2 is also passed to node 1 (Figure 6C). At this point, new embeddings for node 1 are generated using not only its own information, but also that propagated from its 1^st^ and 2^nd^ order neighbors. Three graph convolution blocks are employed in our model because we found empirically that adding one more block does not improve the performance anymore. Further, there is no point of using more than four blocks because the diameter of the cancer-specific graph is 5, so no new information is propagated beyond 4^th^ order neighbors.

**Figure 6.**
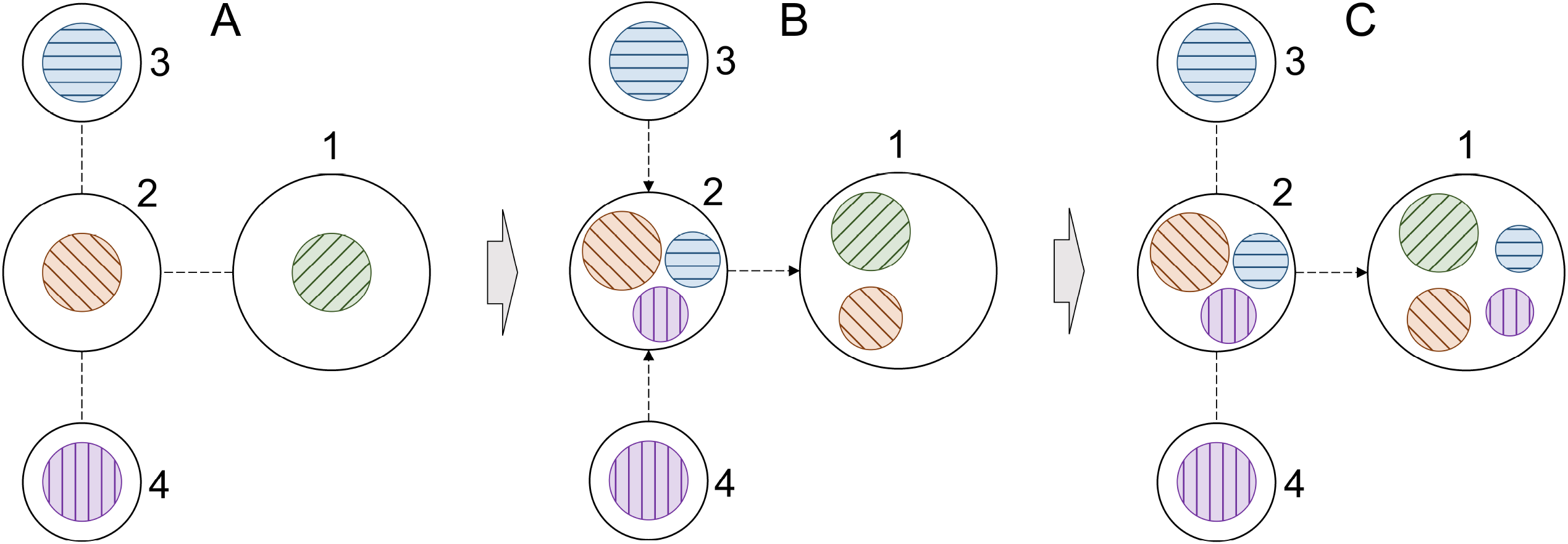
Schematic of information propagation in a graph. **(A)** A simple 4-node graph, in which each node contains its own information. The information is color coded, node 1 – green, node 2 – orange, node 3 – blue, and node 4 – purple. (B) The distribution of information within the graph after the first propagation step. (C) The distribution of information within the graph after the second propagation step. Only the information propagation to node 1 is illustrated in order to demonstrate how it receives information from higher order neighbors.

### Graph information extraction

Once all embeddings are generated, the information on the entire graph can be extracted with a readout mechanism to predict the final drug response (Figure 5C). Standard readout techniques, such as global pooling, are unsuitable for our model comprising multiple graph convolutional blocks and learning from highly heterogeneous input graphs. In GraphGR, node embeddings generated by consecutive graph convolutional blocks contain distinct information. Therefore, a jumping knowledge network (JK-Net) is employed to exploit all information collected from different blocks. JK-Net was specifically developed to efficiently integrate the output from different layers into a single representation [53]. It is based on the concept of an influence radius corresponding to the radius of neighbors whose output is to be aggregated. The selection of an optimal radius is crucial because large radii may cause too much averaging and small radii may result in an insufficient information aggregation. JK-Net learns the effective neighborhood size for each layer in order to generate the best representation of the entire graph.

Global pooling of the embeddings of all nodes is appropriate only for homogeneous networks. In contrast, cancer-specific networks are highly heterogeneous comprising nodes of varying importance to one another and to the overall graph. Therefore, we added a mechanism to emphasize on important nodes rather than treating all nodes equally. Although such techniques have successfully been used in the CNN and the RNN [54], unlike images or text, graphs are orderless, i.e. an image does not remain the same if pixels are rearranged, while a graph remains the same if nodes are reordered. To account for the lack of order in graphs, we added a Set2Set layer converting a set to another set [55]. This model employs a set of LSTMs recursively combining the state of the previous processing step with the current embeddings to generate attentions. These attentions and embeddings form new states for the next processing step. By using Set2Set, we ensure that any permutation performed on the original vector do not affect the final read vector. The information summarized by JK-Net and Set2Set for the entire graph is then passed to a set of fully connected layers for the final prediction, which is the effect of pharmacotherapy on cancer cell growth.

### Performance of GraphGR compared to other methods

In order to properly evaluate the generalizability of GraphGR, we performed a cross-validation at the tissue level. The entire dataset was first divided into nine groups of different tissues, digestive system, respiratory system, haematopoietic and lymphoid tissue, breast tissue, female reproductive system, skin, nervous system, excretory system, and others. Next, we conducted a 9-fold cross-validation, each time using cancer cell lines from one tissue as a validation set while the remaining cancer cell lines were used for model training. Since cell lines collected from different tissues have different gene expression patterns, this cross-validation scheme eliminates the overlap between training and validation data because the reduced graphs have different topologies. In addition, there is also a desired variability in feature matrices on account on different gene-disease associations which depend on the cell line and tissue type. Essentially, each fold has entirely different training and validation data. Figure 7 shows a crossvalidated ROC plot for GraphGR compared to other methods. Indeed, GraphGR not only gives the highest mean ±standard deviation area under the curve (AUC) of 0.83 ±0.03, but the AUC values do not vary significantly for different tissues, digestive system (an AUC of 0.86), respiratory system (an AUC of 0.79), haematopoietic and lymphoid tissue (an AUC of 0.79), breast tissue (an AUC of 0.84), female reproductive system (an AUC of 0.88), skin (an AUC of 0.82), nervous system (an AUC of 0.83), excretory system (an AUC of 0.84), and others (an AUC of 0.78).

**Figure 7.**
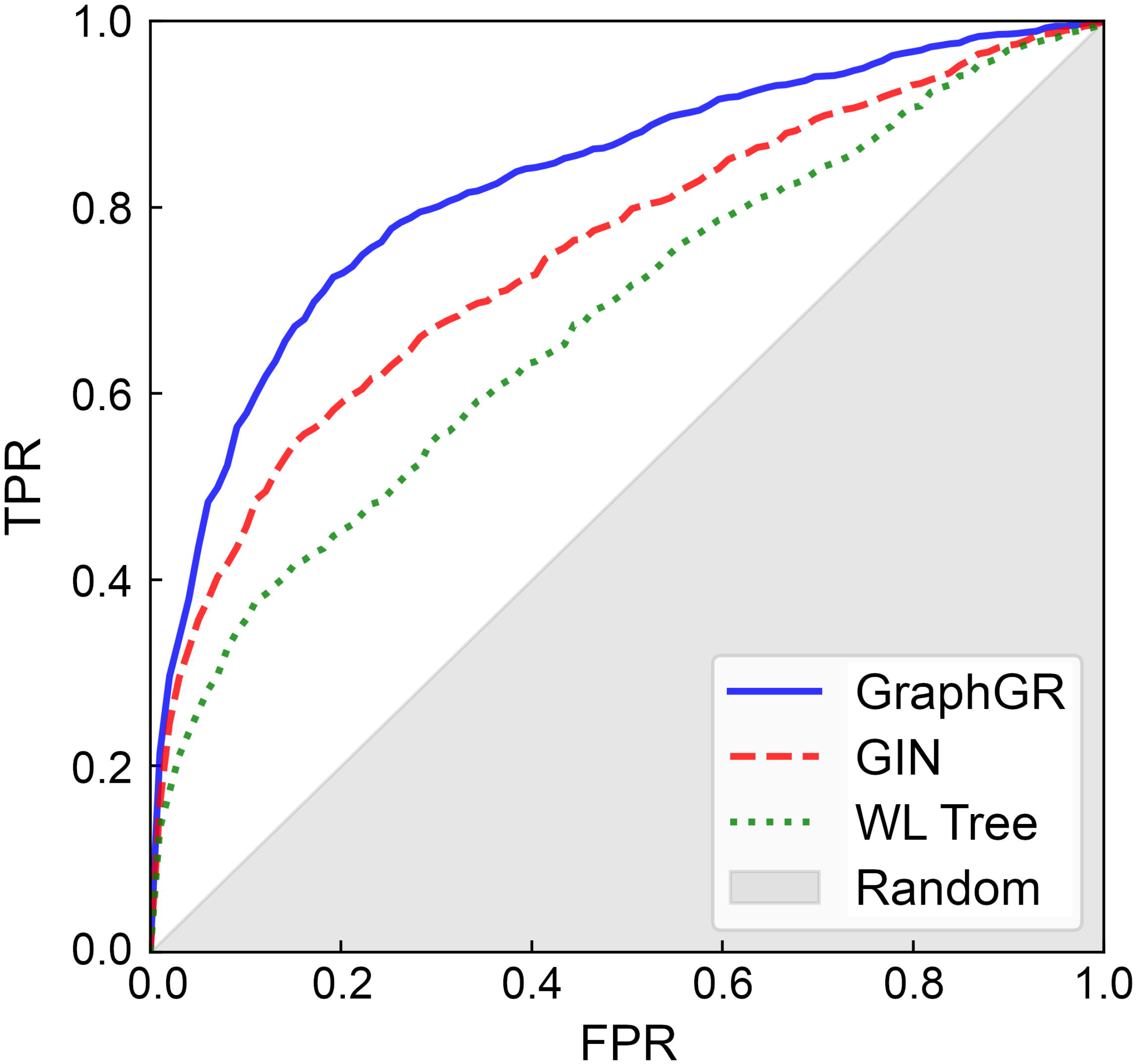
Performance of algorithms to predict the response of cancer cell lines to drugs. The performance of each method is cross-validated at the tissue level. GraphGR (solid blue line) is compared to the graph isomorphism network (GIN, dashed red line) utilizing equal propagation, and WL Tree (dotted green line) employing the Weisfeiler-Lehman graph kernel. TPR is the true positive rate, FPR is the false positive rate, and the gray area corresponds to the performance of a random predictor.

Removing the attention mechanism, which detects important nodes and puts more weight on them (labeled as GIN in Figure 7), decreases the AUC to 0.75 ±0.04 demonstrating that the propagation attention is an important component of GraphGR. Further, the performance of GraphGR is compared to that of the Weisfeiler-Lehman (WL) Tree, a widely adopted graph kernel method for graph machine learning [56]. This algorithm utilizes kernel functions and the WL graph isomorphism test to iteratively generate new labels for nodes and new representations for graphs. Not only the AUC for WL Tree of 0.68 ±0.03 is lower than that for GraphGR, but since WL Tree processes one graph at a time, its runtimes are much longer than those for GraphGR featuring batch processing. Finally, we compare GraphGR with a traditional method to identify potentially effective treatments based on matching the oppose gene signatures. We could conduct this analysis for 635 combinations of 11 cell lines and 22 drugs, for which the perturbational profiles in cancer cell lines are available in the Next Generation Connectivity Map [57]. Using this method yields an AUC of only 0.46 showing that the alteration of gene expression alone cannot reliably predict the drug response for cancer cell lines. Overall, these results demonstrate that GraphGR clearly outperforms other deep learning, graph kernel, and traditional approaches.

### GraphGR predictions validated by literature

We performed independent testing of GraphGR on several cases not present in the LINCS growth rate inhibition dataset, thus not included in the training and validation sets. Each new prediction is supported by evidence found in the biomedical literature. The first case is motesanib, an anthranilamide inhibitor of vascular endothelial growth factor receptors (VEGFR) with IC_50_ values of 2 nM (VEGFR1), 3 nM (VEGFR2), and 6 nM (VEGFR3) [58]. Though VEGFR kinases are its primary targets, motesanib also inhibits the activity of platelet derived growth factor receptor beta (PDGFRB) at an IC_50_ of 84 nM, mast/stem cell growth factor receptor Kit (c-KIT) at an IC_50_ of 8 nM, and tyrosine-protein kinase receptor Ret (c-RET) at an IC_50_ of 59 nM [59]. This drug has been tested alone and in combination with chemotherapy in human non-smallcell lung cancer xenograft models created by injecting NCI-H358, NCI-H1299, NCI-H1650, A549, and Calu-6 cancer cell lines subcutaneously into mice. Tested against A549 at three different concentrations, 7.5, 25, and 75 mg/kg b.i.d, motesanib inhibited the tumor growth by 45%, 84%, and 107%, respectively. GraphGR estimated a high probability of 0.82 for the growth inhibition of A549 cell line by motesanib. Further, the tumor growth of Calu-6 xenograft was inhibited by 66% at the highest tested dose of motesanib [60]. Encouragingly, the probability that motesanib inhibits the growth of Calu-3 cell line reported by GraphGR is as high as 0.97. Note that according to the Cellosaurus [61], Calu-3 (originated from 25 years old male) and Calu-6 (originated from 61 years old female) are closely related lung adenocarcinoma cell lines.

Motesanib also has antitumor activity against breast cancer [58]. Its primary targets, VEGFR proteins, are angiogenic factors that modulate processes playing important roles in the development and progression of breast cancer [62]. Motesanib was tested against MCF-7, MDA-MB-231, and Cal-51 xenografts of breast cancer. It inhibited MCF-7 tumor growth by 44% at a concentration of 25 mg/kg and by 65% at a concentration of 75 mg/kg. Further, motesanib inhibited MDA-MB-231 tumor growth by 64% at the highest concentration. Cal-51 tumor growth was also reduced by 38%, 74% and 81% when the drug was administered at 7.5 mg/kg, 25 mg/kg and 75 mg/kg, respectively [62]. GraphGR estimated that the probabilities of inhibiting the growth of MCF-7, MDA-MB-231, and Cal-51 breast cancer cell lines are 0.88, 0.95, and 0.93, respectively.

Pazopanib inhibits intracellular tyrosine kinases, PDGFRA with an IC_50_ of 73 nM, PDGFRB with an IC_50_ of 215 nM, VEGFR1 with an IC_50_ of 7 nM, VEGFR2 with an IC_50_ of 15 nM, and VEGFR3 with an IC_50_ of 2 nM [63]. It exhibits antiangiogenic properties and is used to treat renal cell carcinoma (RCC) [64]. Pazopanib was tested in 8 human RCC cell lines, 769-P, 786-0, HRC-24, HRC-31, HRC-45, HRC-78, RCC-26B, and SK-45, showing a varying degree of antiproliferative activities [65]. For instance, it reduces the proliferation of 786-O cell lines by 50% at 100 μM. According to GraphGR, the probability of inhibition of the 786-O cell line growth by pazopanib is 0.76. Pazopanib was also tested alone and in combination with topotecan against anaplastic thyroid cancer (cell line 8305C) [66], one of the most aggressive, but rare forms of thyroid cancer. 72 hours after the treatment with pazopanib, the proliferation of 8305C cell line was inhibited at an IC_50_ of 25 μM ±3.2. According to the Cellosaurus, 8305C (originated from 67 years old female) and 8505C (originated from 78 years old female) cell lines are closely related anaplastic thyroid cancers and GraphGR estimated that pazopanib inhibits the growth of 8505C with a high probability of 0.93.

Lestaurtinib is a multitargeted kinase inhibitor structurally related to staurosporine [67]. It inhibits fms-like tyrosine kinase 3 (FLT3) with an IC_50_ of 2 to 3 nM [68], Janus kinase 2 (JAK2) with an IC_50_ of 1 nM [69], and tropomyosin receptor kinases (Trk) with an IC_50_ of 100 nM [70]. Human pancreatic ductal adenocarcinoma (PDAC) shows an aberrant expression of neurotrophin and its associated Trk receptors [71]. After the drug was administered at 10 mg/kg b.i.d into a mouse model created by subcutaneously injecting a PDAC cell line Panc1, the growth of the xenograft showed a significant decrease with a *p*-value of <0.01 [71]. GraphGR predicted with a high probability of 0.98 that lestaurtinib inhibits the growth of Panc 04.03, which is a closely related PDAC cell line.

## Methods

### Growth rate inhibition data

Although IC_50_ and E_max_ are conventional measures of the effect of drugs on cell proliferation, these metrics solely dependent on the number of cell divisions throughout the treatment time. On the other hand, more recent drug response metrics, GR50 and GR_max_, quantify the proliferation with the value of growth rate inhibition (GR) based on time course and endpoint assays [72]. GR50 is the concentration of a drug at which GR is 0.5, whereas GR_max_ is the maximum measured GR value. GR_max_ values range from −1 to 1 with negative values corresponding to the cytotoxic response and positive values corresponding to the cytostatic response. In this study, we employ six LINCS-Dose-Response datasets, Broad-HMS LINCS Joint Project, LINCS MCF10A Common Project, HMS LINCS Seeding Density Project, MEP-HMS LINCS Joint Project, Genentech Cell Line Screening Initiative, and Cancer Therapeutics Response Portal [72]. The original dataset contains 632 cell lines from different cancer tissues and 795 small molecules tested against those cancer cell lines, totaling 83,162 combinations. After removing those cases having either GR50 values set to infinity or multiple GR50 values for a particular cell line-drug combination, the final dataset comprises 359 cell lines, 57 drugs, and 6,167 cell line-drug combinations.

### Protein-protein interaction network

Protein-protein interaction data were acquired from the STRING database [73] that comprises 19,354 human proteins forming 11,355,804 interactions including those identified experimentally as well as predicted computationally. Each interaction is assigned a confidence score ranging from 150 for low-confidence to 999 for high-confidence interactions. A human PPI network was constructed from confident interactions with a score of ≥500. Single proteins disconnected from the main network and those forming small isolated networks were removed, resulting in the final PPI network containing 19,144 proteins and 685,198 interactions.

### Differential gene expression data

Original gene expression data are available from the Cancer Cell Line Encyclopedia (CCLE) project comprising a detailed genetic characterization of a large number of cancer cell lines [27]. We obtained the curated CCLE data from Harmonizome, a comprehensive repository of processed genomics, proteomics, epigenomics, transcriptomics, and metabolomics data [74]. The differential gene expression dataset contains 18,022 genes, 1035 cancer cell lines, and 749,551 gene-cell associations categorized as down-, up-, and normally regulated with respect to the expression level in healthy cells.

### Kinase inhibitor profiling

Inhibitor profiling refers to a large-scale experimental measurement of the activity of an inhibitor against a panel of target proteins. Kinase inhibitor profiling data used in this study were collected and curated by Team-SKI [75]. The activity is reported as a pIC_50_ value, which is the negative logarithm of the half-maximal inhibitory concentration (IC_50_) measuring the potency of a compound in inhibiting a specific biochemical function. The cutoff for pIC_50_ values was set at 6.3 corresponding to the IC_50_ of 500 nM. The Team-SKI dataset contains 49,348 small molecules tested against 411 protein kinases.

### Disease-gene associations

Disease-gene association scores were obtained from two sources, DISEASES [76] and DisGeNET [77]. The DISEASES database integrates evidence on associations collected through automatic text mining, manually curated from biomedical literature, cancer mutation data, and genomewide association studies. It contains 8,330 diseases and 20,715 genes with association scores ranging from 1 to 10. The DisGeNET database provides information on associations pulled from various repositories including Mendelian, complex and environmental diseases, and integrated using gene/disease vocabulary mapping and the DisGeNET ontology. It comprises 24,166 diseases and 17,545 genes with association scores ranging from 0.01 to 1.

### Data integration

Data integration was performed by mapping gene expression, inhibitor profiling, and diseasegene associations directly onto the PPI network. Gene expression data contains normalized scores for 18,022 genes, whereas the PPI network comprises 19,144 proteins. The missing values for nodes in the graph were replaced by median scores calculated over 1^st^ order neighbors. The kinase component of the PPI network was determined by running Basic Local Alignment Search Tool (BLAST) [78] against known human kinases [79]. Using a similarity threshold of 95% identified 508 kinases. pIC_50_ values are available from the Team-SKI database for 411 kinase proteins in our network and 29 small molecules that are also present in the growth rate inhibition data from LINCS. The Disease Ontology ID (DOID) and the Concept ID for each cancer cell line were identified with the Cellosaurus resource portal [61], and mapped to, respectively, DISEASES and DisGeNET databases in order to assign disease-gene association scores to nodes in the PPI network. After data integration and removing cases that could not be mapped, the final dataset contains annotated graphs for 3,549 combinations of 359 cell lines and 29 drugs.

### Graph partitioning

The latest release of the GO database contains 29,698 BP, 11,147 MF, and 4,201 CC ontology terms [50]. Each DAG has its own hierarchy maintained by the domain-centric Gene Ontology (dcGO). According to the dcGO database, the numbers of level-1 children nodes are 30, 15, and 22 for BP, MF, and CC, respectively [52]. Since our goal was to assign nodes to biological pathways, all against all semantic similarities between GO terms of PPI network nodes were calculated with the GOGO software based on the BP-DAG topology. The matrix of GOGO scores was then subjected to a hierarchical clustering analysis, which builds nested clusters by successively merging and splitting clusters [80]. There are two types of hierarchical clustering, agglomerative (bottom-up) and divisive (top-down). Agglomerative methods merge observations as moving up the hierarchy while divisive methods split the observations as moving down the hierarchy. Here, we employed the agglomerative HCA rather than other commonly used clustering methods, such as K-means [81], DBSCAN [42], and affinity propagation [43] because this technique is the most appropriate for non-Euclidean GOGO semantic similarities and we found empirically that the clustering results represent well different levels of biological process in GO.

### Graph density

Graph density, *ρ_G_*, is defined as the ratio of the number of edges and the number of possible edges. For an undirected graph, it is calculated as

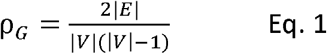

where |*E*| is the number of edges and |*V*| is the number of nodes. Since the information is propagated more efficiently across dense graphs, increasing the graph density is generally beneficial for graph-based machine learning algorithms, such as GNN.

### Graph-feature entropy

The graph reduction procedure basically compresses the information in a graph. However, as all lossy compression schemes, some information will be lost during the reduction. In order to measure how much meaningful information is lost or gained as opposed to the loss/gain of the redundant or irrelevant information, we recently introduced a concept of the graph-feature entropy. Briefly, the graph-feature entropy S’ of a graph *G, S_G_* is based on the Shannon entropy [82] defined as

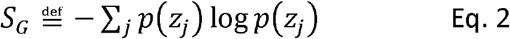

where *Z_j_* is the column vector of the feature matrix filtered by the graph Laplacian, which corresponds to the *j*-th feature of all nodes in the graph. Using graph Laplacian filtered features effectively combines both feature and topological information of a graph providing a useful measure of the information content in featured graphs.

Using the Shannon entropy of features and the graph-feature-entropy, we calculate the information gain/loss after reduction, δ, as

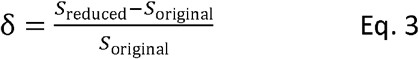

where *S_original_* is the entropy (either feature-only or graph-feature) of the original graph and *S_reduced_* is the corresponding entropy of the reduced graph.

### Information propagation

The most widely adopted propagation protocol transmit the information equally without considering the importance of a node to its neighbors and to the graph. This protocol can be expressed as

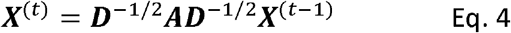

where *t* is the propagation step, ***D*** is the degree matrix of the adjacency matrix ***A***, and ***X***^(*t*)^ represents embeddings at the propagation step ***X***^(0)^. Note that the original node features can be denoted as the 0-th propagation step, It is obvious that not all nodes have the same importance to their neighbors. For instance, many non-kinases in our dataset contain no useful information because these proteins are normally expressed, have no association with a disease, and are not targets for inhibitors. The information propagating from such proteins should be less important compared to the information coming from kinases and other proteins differentially expressed and having high disease associations. On that account, we added a propagation attention mechanism to increase the importance of these nodes. Specifically, we implemented a mechanism to learn a dynamic and adaptive summary of the local neighborhood, which operates only in the feature space [83]. The attention from node *i* to node *j, γ_i,j_*, is defined as

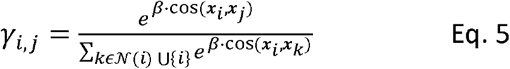

where 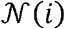 denotes the neighbor nodes of *i* and *β* is a trainable parameter. Essentially, the attention is the softmax of feature cosine similarities between the centering nodes and its neighbors. By utilizing the attention mechanism, the original propagation matrix calculated from the degree and adjacency matrices (see Eq. 4) can be replaced with a new propagation matrix **Γ**, which adaptively adjusts propagation weights based on neighbor features. This new propagation scheme addressing the problem of equal weights can be expressed as

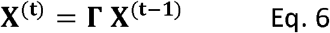

where each entry of the propagation matrix **Γ** is calculated using Eq. 5.

### Node embeddings

After the information is propagated, the embeddings of each node need to be updated. Many techniques are available to generate node embeddings, and each has its advantages and disadvantages. Based on a series of preliminary experiments, we decided to implement a model inspired by the graph isomorphism network (GIN) [84]. The GIN offers an exceptional performance and has a relatively simple structure, which is important for our model because even after reduction, the cancer input data are much larger than typical datasets used by researchers in other fields. Briefly, the GIN transforms the graph isomorphism to the context of deep learning. Nonetheless, it employs a rather basic propagation scheme summing up features from all neighbor nodes. In order to further increase the performance, we replaced this simple propagation step with the attention-based propagation scheme (see Eq. 6). Combining the GIN update protocol with the propagation attention results in a very efficient graph convolution block expressed as

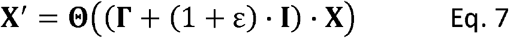

where **Θ** denotes a neural network, **Γ** is the propagation matrix calculated using Eq. 5 and ε is a trainable parameter.

### Graph readout mechanism

GraphGR employs JK-Net followed by a Set2Set model to generate a global representation of the input graph from the node-wise information. JK-Net exploits varying influence radii of different layers to learn the best representation of the entire graph. This model can integrate outputs from individual graph convolutional blocks with three strategies, the concatenation, the max-pooling, and the LSTM-attention. Considering the size of our data, we decided to employ the max-pooling strategy since it does not introduce any additional hyperparameters. This particular strategy performs a feature-wise max-pooling with lower layers favoring the local information and higher layers mostly containing the global graph information. With the max-pooling scheme, JK-Net automatically selects the most informative neighborhood size for each feature coordinate. With the information from different layers aggregated, we adopted the Set2Set model [55] as a final attention-based readout mechanism. A conventional method to simply flatten all embeddings is unsuitable for orderless graphs, which require a premutation-invariant readout mechanism instead. Set2Set comprises three blocks, a reading block, a process block, and a write block. In GraphGR, the reading block generating embeddings for each item in the set is replaced by JK-Net aggregating information from multiple graph convolutional blocks. The process block is an LSTM that reads the embeddings and state generated from the previous processing step, and outputs a new hidden state. Finally, the write block also is an LSTM, which takes the hidden state as a context to generate the attention for each item in the set. Subsequently, the attention vector is combined with the embedding matrix using a weighted summation to generate new, permutation-invariant embeddings.

### Other methods to predict drug response

The Weisfeiler-Lehman (WL) graph kernel, also known as WL Tree, was selected for comparative benchmarks against GraphGR because this method can use the same input data, which are the reduced representations of cancer-specific networks. WL Tree is a widely adopted kernel method for learning from large graphs with discrete node labels [56]. A key component of this algorithm is a rapid feature extraction employing the WL test of isomorphism on graphs. Briefly, WL Tree iteratively maps the original node labels to new node labels based on the topological structure of the graph. The new node labels are then compressed and integrated with the original node labels in order to form a vector representation for the graph. The implementation of WL Tree used in our study was demonstrated to outperform other graph kernels on several graph classification benchmark datasets in terms of accuracy and runtime.

Comparing gene signatures constructed for drug-treated and untreated disease cell lines is a traditional method to find potentially effective therapeutics. The comparison is based on the cosine distance (COS) between two types of gene expression signatures

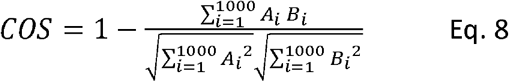

where *A* is the gene signature of a cancer cell line against a healthy cell line and *B* is the gene signature of the same cancer cell line before and after drug treatment. Each gene signature comprises 1000 landmark genes whose expression levels were measured with the L1000 technology. Signatures *A* are constructed using level-5 moderated Z-scores (MODZ) from L1000CDS^2^ [85], whereas signatures *B* are generated based on changes in gene expression levels measured at six drug concentrations, 0.04, 0.12, 0.37, 1.11, 3.33, 10 μM, available from the Next Generation Connectivity Map [57]. COS values range from 0 for exactly the same gene signatures to 2 for exactly opposite gene signatures. A gene signature *B* for the drug treatment opposing a gene signature *A* for the disease indicates that the drug may reverse expression changes related the disease state. We calculated six COS values for all drug concentrations and then selected the longest distance as a metric to predict the anticancer drug response.

## Discussion

In this study, we developed GraphGR, a graph neural network model to predict the growth rate of a cancer cell line after drug treatment. GraphGR learns from cancer-specific networks constructed by integrating multiple heterogeneous data, including protein-protein interactions, differential gene expression, gene-disease associations, and kinase inhibitor profiling. In order to improve the effectiveness of machine learning, a graph reduction protocol was devised to convert initially information-sparse biological networks to information-rich graphs preserving the biological knowledge. In contrast to the original PPI networks that share exactly the same topology, the reduced graphs provide a diverse set of graph topologies across different cell lines. In addition, because of the much smaller size compared to full PPI networks, the reduced graphs can be used with any deep learning model, even those that are computationally expensive. Overall, the reduction procedure not only enables a more efficient learning process, but it also improves the performance of GraphGR on account of the more diverse data.

GraphGR is more advanced than many deep learning techniques operating in the Euclidean space [36], because it extracts knowledge directly from biological networks, which provide a more adequate representation of complex diseases such as cancer. Further, we implemented a sophisticated attention mechanism to more efficiently propagate information from the most important nodes in the graph when generating node embedding. Attention mechanisms assigning trainable weights to nodes during information propagation are used to improve not only the classification performance [86, 87], but also the capability to generalize to larger, more complex, and noisy graphs [88, 89]. In our case, this technique allows the GNN model to direct more attention to kinase nodes since many of them contain valuable information on differential gene expression and the level of inhibition by various compounds across different cancer cell lines. As a result, the GNN achieves a better performance, especially against highly heterogeneous networks, such as cancer-specific networks constructed in this study.

In order to evaluate the performance of GraphGR, we conducted a cross-validation at the tissue level by removing from model training all cell lines originating in a particular tissue and then analyzing the accuracy for these cell lines. We put a special attention to design a proper benchmarking protocol since in the context of predictive models, misunderstanding cross-validation very often yields an impressive, yet grossly overestimated predictor performance [90]. Numerous examples of exaggerated results in biomedical studies due to a problematic cross-validation include cancer prediction [91], the prediction of cancer cell line sensitivity and compound potency [92], the identification of drug-target interactions [93], the prediction of optimal drug therapies [94], the estimation of drug-target binding affinities [95], and virtual screening [96]. Since multiple instances in our dataset share cell lines originating from the same tissue, employing cross-validation at the tissue level is critical because splitting the dataset randomly into folds would cause training and validation instances to have a significant overlap with respect to graph topology as well as certain features such as gene expression and gene-disease associations.

Comparative benchmarking calculations demonstrate that GraphGR not only outperforms other deep learning methods, graph kernel models, and traditional algorithms, but it also generalizes well to unseen data. For independent testing, we selected several kinase inhibitors with no growth rate inhibition data in LINCS, therefore not included in the training and validation sets. The effect of these testing molecules on the growth of several cell lines predicted by GraphGR was validated against the biomedical literature. Encouragingly, those inhibitors assigned high probabilities to reduce the proliferation of certain cell lines have been reported to exhibit anticancer activities in independent experiments. Overall, these results indicate that GraphGR is suitable for a broad range of applications involving a variety of cancer cell lines and inhibitors.

## Availability

GraphGR is freely available to the academic community at https://github.com/pulimeng/GraphGR.

## Acknowledgements

Portions of this research were conducted with computing resources provided by Louisiana State University.

## References

1. Knox, S.S., From ‘omics’ to complex disease: a systems biology approach to gene-environment interactions in cancer. Cancer Cell Int, 2010. 10: p. 11.

2. Cicenas, J., et al., Kinases and cancer. Cancers (Basel), 2018. 10(3).

3. Bhullar, K.S., et al., Kinase-targeted cancer therapies: progress, challenges and future directions. Mol Cancer, 2018. 17(1): p. 48.

4. McDermott, U. and J. Settleman, Personalized cancer therapy with selective kinase inhibitors: an emerging paradigm in medical oncology. J Clin Oncol, 2009. 27(33): p. 5650–9.

5. Brylinski, M. and J. Skolnick, Comprehensive structural and functional characterization of the human kinome by protein structure modeling and ligand virtual screening. J Chem Inf Model, 2010. 50(10): p. 1839–54.

6. Brylinski, M. and J. Skolnick, Cross-reactivity virtual profiling of the human kinome by X-react(KIN): a chemical systems biology approach. Mol Pharm, 2010. 7(6): p. 2324–33.

7. Davis, M.I., et al., Comprehensive analysis of kinase inhibitor selectivity. Nat Biotechnol, 2011. 29(11): p. 1046–51.

8. Hartmann, J.T., et al., Tyrosine kinase inhibitors - a review on pharmacology, metabolism and side effects. Curr Drug Metab, 2009. 10(5): p. 470–81.

9. Yang, X., et al., Kinase inhibition-related adverse events predicted from in vitro kinome and clinical trial data. J Biomed Inform, 2010. 43(3): p. 376–84.

10. Gujral, T.S., L. Peshkin, and M.W. Kirschner, Exploiting polypharmacology for drug target deconvolution. Proc Natl Acad Sci USA, 2014. 111(13): p. 5048–53.

11. Knight, Z.A., H. Lin, and K.M. Shokat, Targeting the cancer kinome through polypharmacology. Nat Rev Cancer, 2010. 10(2): p. 130–7.

12. Ma, X., X. Lv, and J. Zhang, Exploiting polypharmacology for improving therapeutic outcome of kinase inhibitors (KIs): An update of recent medicinal chemistry efforts. Eur J Med Chem, 2018. 143: p. 449–463.

13. Fedorov, O., F.H. Niesen, and S. Knapp, Kinase inhibitor selectivity profiling using differential scanning fluorimetry. Methods Mol Biol, 2012. 795: p. 109–18.

14. Schirle, M., et al., Kinase inhibitor profiling using chemoproteomics. Methods Mol Biol, 2012. 795: p. 161–77.

15. Duong-Ly, K.C., et al., Kinase Inhibitor Profiling Reveals Unexpected Opportunities to Inhibit Disease-Associated Mutant Kinases. Cell Rep, 2016.14(4): p. 772–781.

16. Jacoby, E., et al., Extending kinome coverage by analysis of kinase inhibitor broad profiling data. Drug Discov Today, 2015. 20(6): p. 652–8.

17. Miduturu, C.V., et al., High-throughput kinase profiling: a more efficient approach toward the discovery of new kinase inhibitors. Chem Biol, 2011. 18(7): p. 868–79.

18. Garnis, C., T.P. Buys, and W.L. Lam, Genetic alteration and gene expression modulation during cancer progression. Mol Cancer, 2004. 3: p. 9.

19. Hanahan, D. and R.A. Weinberg, The hallmarks of cancer. Cell, 2000. 100(1): p. 57–70.

20. Lo, K.C., et al., Identification of genes involved in squamous cell carcinoma of the lung using synchronized data from DNA copy number and transcript expression profiling analysis. Lung Cancer, 2008. 59(3): p. 315–31.

21. Hahn, W.C. and R.A. Weinberg, Rules for making human tumor cells. N Engl J Med, 2002. 347(20): p. 1593–603.

22. Ismail, R.S., et al., Differential gene expression between normal and tumor-derived ovarian epithelial cells. Cancer Res, 2000. 60(23): p. 6744–9.

23. Liang, P. and A.B. Pardee, Analysing differential gene expression in cancer. Nat Rev Cancer, 2003. 3(11): p. 869–76.

24. Chen, S., B. Zhu, and L. Yu, In silico comparison of gene expression levels in ten human tumor types reveals candidate genes associated with carcinogenesis. Cytogenet Genome Res, 2006. 112(1-2): p. 53–9.

25. Deng, J.L., Y.H. Xu, and G. Wang, Identification of Potential Crucial Genes and Key Pathways in Breast Cancer Using Bioinformatic Analysis. Front Genet, 2019.10: p. 695.

26. Xue, J.M., et al., Comprehensive Analysis of Differential Gene Expression to Identify Common Gene Signatures in Multiple Cancers. Med Sci Monit, 2020. 26: p. e919953.

27. Chang, Y., et al., Cancer drug response profile scan (CDRscan): A deep learning model that predicts drug effectiveness from cancer genomic signature. Sci Rep, 2018. 8(1): p. 8857.

28. Wang, L., et al., Improved anticancer drug response prediction in cell lines using matrix factorization with similarity regularization. BMC Cancer, 2017.17(1): p. 513.

29. Gyurko, D.M., et al., Adaptation and learning of molecular networks as a description of cancer development at the systems-level: potential use in anti-cancer therapies. Semin Cancer Biol, 2013. 23(4): p. 262–9.

30. Klinke, D.J., 2nd, Signal transduction networks in cancer: guantitative parameters influence network topology. Cancer Res, 2010. 70(5): p. 1773–82.

31. Liu, Y., et al., A multiscale computational approach to dissect early events in the Erb family receptor mediated activation, differential signaling, and relevance to oncogenic transformations. Ann Biomed Eng, 2007. 35(6): p. 1012–25.

32. Chen, C., et al., Construction and analysis of protein-protein interaction networks based on proteomics data of prostate cancer. Int J Mol Med, 2016. 37(6): p. 1576–86.

33. Guda, P., S.V. Chittur, and C. Guda, Comparative analysis of protein-protein interactions in cancer-associated genes. Genomics Proteomics Bioinformatics, 2009. 7(1-2): p. 25–36.

34. Kanhaiya, K., et al., Controlling Directed Protein Interaction Networks in Cancer. Sci Rep, 2017. 7(1): p. 10327.

35. Erten, S., G. Bebek, and M. Koyuturk, Vavien: an algorithm for prioritizing candidate disease genes based on topological similarity of proteins in interaction networks. J Comput Biol, 2011. 18(11): p. 1561–74.

36. Bronstein, M.M., et al., Geometric deep learning: Going beyond Euclidean data. IEEE Signal Processing Magazine, 2017. 34(4): p. 18–42.

37. Scarselli, F., et al., The graph neural network model. IEEE Trans Neural Netw, 2009. 20(1): p. 61–80.

38. Kipf, T.N. and M. Welling, Semi-supervised classification with Graph Convolutional Networks. arXiv, 2016: p. 1609.02907.

39. Kipf, T.N. and M. Welling, Variational graph auto-encoders. arXiv, 2016: p. 1611.07308.

40. Li, Y., et al., Gated graph sequence neural networks. arXiv, 2015: p. 1511.05493.

41. Hamilton, W.L., R. Ying, and J. Leskovec, Inductive representation learning on large graphs. arXiv, 2018: p. 1706.02216.

42. Bacciu, D., F. Errica, and A. Micheli, Contextual Graph Markov Model: A deep and generative approach to graph processing. arXiv, 2018: p. 1805.10636.

43. Chen, J., T. Ma, and C. Xiao, FastGCN: Fast learning with graph convolutional networks via importance sampling. arXiv, 2018: p. 1801.10247.

44. Liang, X., et al., Semantic object parsing with graph LSTM. arXiv, 2016: p. 1603.07063.

45. Golub, G.H. and C.F. Van Loan, Matrix computations. 3rd ed. Johns Hopkins studies in the mathematical sciences. 1996, Baltimore: Johns Hopkins University Press, xxvii, 694 p.

46. Rosen, K.H., Discrete mathematics and its applications. 7th ed. 2012, New York: McGraw-Hill.

47. Gross, J.L. and J. Yellen, Graph theory and its applications. The CRC Press series on discrete mathematics and its applications. 1999, Boca Raton, Fla.: CRC Press. 585 p.

48. West, D.B., Introduction to graph theory. 2nd ed. 2001, Upper Saddle River, N.J.: Prentice Hall, xix, 588 p.

49. Zhao, C. and Z. Wang, GOGO: An improved algorithm to measure the semantic similarity between gene ontology terms. Sci Rep, 2018. 8(1): p. 15107.

50. Ashburner, M., et al., Gene ontology: tool for the unification of biology. The Gene Ontology Consortium. Nat Genet, 2000. 25(1): p. 25–9.

51. Sharan, R., I. Ulitsky, and R. Shamir, Network-based prediction of protein function. Mol Syst Biol, 2007. 3: p. 88.

52. Fang, H. and J. Gough, dcGO: database of domain-centric ontologies on functions, phenotypes, diseases and more. Nucleic Acids Research, 2013. 41(D1): p. D536–D544.

53. Xu, K., et al., Representation learning on graphs with jumping knowledge networks. arXiv e-prints, 2018: p. arXiv:1806.03536.

54. Vaswani, A., et al., Attention Is All You Need. arXiv e-prints, 2017: p. arXiv:1706.03762.

55. Vinyals, O., S. Bengio, and M. Kudlur, Order matters: Sequence to sequence for sets. arXiv e-prints, 2015: p. arXiv:1511.06391.

56. Shervashidze, N., et al., Weisfeiler-Lehman graph kernels. J Mach Learn Res, 2011. 12: p. 2539–2561.

57. Subramanian, A., et al., A next generation Connectivity Map: L1000 platform and the first 1,000,000profiles. Cell, 2017. 171(6): p. 1437–1452 e17.

58. Musumeci, F., et al., Vascular endothelial growth factor (VEGF) receptors: drugs and new inhibitors. J Med Chem, 2012. 55(24): p. 10797–822.

59. Polverino, A., et al., AMG 706, an oral, multikinase inhibitor that selectively targets vascular endothelial growth factor, platelet-derived growth factor, and kit receptors, potently inhibits angiogenesis and induces regression in tumor xenografts. Cancer Res, 2006. 66(17): p. 8715–21.

60. Coxon, A., et al., Antitumor activity of motesanib alone and in combination with cisplatin or docetaxel in multiple human non-small-cell lung cancer xenograft models. Mol Cancer, 2012. 11: p. 70.

61. Bairoch, A., The Cellosaurus, a Cell-Line Knowledge Resource. J Biomol Tech, 2018. 29(2): p. 25–38.

62. Coxon, A., et al., Broad antitumor activity in breast cancer xenografts by motesanib, a highly selective, oral inhibitor of vascular endothelial growth factor, platelet-derived growth factor, and Kit receptors. Clin Cancer Res, 2009. 15(1): p. 110–8.

63. Zhao, H.L., et al., Overview of fundamental study of pazopanib in cancer. Thorac Cancer, 2014. 5(6): p. 487–93.

64. Keisner, S.V. and S.R. Shah, Pazopanib: the newest tyrosine kinase inhibitor for the treatment of advanced or metastatic renal cell carcinoma. Drugs, 2011. 71(4): p. 443–54.

65. Canter, D., et al., Are all multi-targeted tyrosine kinase inhibitors created equal? An in vitro study of sunitinib and pazopanib in renal cell carcinoma cell lines. Can J Urol, 2011. 18(4): p. 5819–25.

66. Di Desidero, T., et al., Effects of pazopanib monotherapy vs. pazopanib and topotecan combination on anaplastic thyroid cancer cells. Front Oncol, 2019. 9: p. 1202.

67. Shabbir, M. and R. Stuart, Lestaurtinib, a multitargeted tyrosine kinase inhibitor: from bench to bedside. Expert Opin Investig Drugs, 2010. 19(3): p. 427–36.

68. Knapper, S., et al., A phase 2 trial of the FLT3 inhibitor lestaurtinib (CEP701) as first-line treatment for older patients with acute myeloid leukemia not considered fit for intensive chemotherapy. Blood, 2006.108(10): p. 3262–70.

69. Hexner, E.O., et al., Lestaurtinib (CEP701) is a JAK2 inhibitor that suppresses JAK2/STAT5 signaling and the proliferation of primary erythroid cells from patients with myeloproliferative disorders. Blood, 2008. 111(12): p. 5663–71.

70. Camoratto, A.M., et al., CEP-751 inhibits TRK receptor tyrosine kinase activity in vitro exhibits anti-tumor activity. Int J Cancer, 1997. 72(4): p. 673–9.

71. Miknyoczki, S.J., et al., The novel Trk receptor tyrosine kinase inhibitor CEP-701 (KT-5555) exhibits antitumor efficacy against human pancreatic carcinoma (Panc1) xenograft growth and in vivo invasiveness. Ann N Y Acad Sci, 1999. 880: p. 252–62.

72. Hafner, M., et al., Growth rate inhibition metrics correct for confbunders in measuring sensitivity to cancer drugs. Nat Methods, 2016. 13(6): p. 521–7.

73. Szklarczyk, D., et al., STRING v11: protein-protein association networks with increased coverage, supporting functional discovery in genome-wide experimental datasets. Nucleic Acids Res, 2019. 47(D1): p. D607–D613.

74. Rouillard, A.D., et al., The harmonizome: a collection of processed datasets gathered to serve and mine knowledge about genes and proteins. Database (Oxford), 2016. 2016.

75. Sorgenfrei, F.A., S. Fulle, and B. Merget, Kinome-wide profiling prediction of small molecules. ChemMedChem, 2018. 13(6): p. 495–499.

76. Pletscher-Frankild, S., et al., DISEASES: text mining and data integration of disease-gene associations. Methods, 2015. 74: p. 83–9.

77. Pinero, J., et al., DisGeNET: a comprehensive platform integrating information on human disease-associated genes and variants. Nucleic Acids Res, 2017. 45(D1): p. D833–D839.

78. Altschul, S.F., et al., Basic local alignment search tool. J Mol Biol, 1990. 215(3): p. 403–10.

79. Manning, G., et al., The protein kinase complement of the human genome. Science, 2002. 298(5600): p. 1912–34.

80. Maimon, O. and L. Rokach, Data mining and knowledge discovery handbook. 2nd ed. 2010, New York: Springer, xx, 1285 p.

81. MacQueen, J.B. Some methods for classification and analysis of multivariate observations, in Proceedings of 5th Berkeley Symposium on Mathematical Statistics and Probability. 1967. Berkeley, CA: University of California Press.

82. Shannon, C.E., A mathematical theory of communication. Bell System Technical Journal, 1948. 27: p. 379–423.

83. Thekumparampil, K.K., et al., Attention-based Graph Neural Network for Semisupervised Learning. arXiv e-prints, 2018: p. arXiv:1803.03735.

84. Xu, K., et al., How Powerful are Graph Neural Networks? arXiv e-prints, 2018: p. arXiv: 1810.00826.

85. Duan, Q., et al., L1000CDS(2): LINCS L1000 characteristic direction signatures search engine. NPJ Syst Biol Appl, 2016. 2.

86. Zhang, S. and L. Xie, Improving attention mechanism in graph neural networks via cardinality preservation. arXiv preprint arXiv:1907.02204, 2019.

87. Zhang, Y., et al., Hyperbolic graph attention network. arXiv preprint arXiv:1912.03046, 2019.

88. Knyazev, B., G.W. Taylor, and M.R. Amer, Understanding attention and generalization in graph neural networks. arXiv preprint arXiv:1905.02850, 2019.

89. Shi, M., et al., Feature-attention graph convolutional networks for noise resilient learning. arXiv preprint arXiv:1912.11755, 2019.

90. Neunhoeffer, M. and S. Sternberg, How cross-validation can go wrong and what to do about it. Political Analysis, 2019. 27(1): p. 101–106.

91. Xiao, Y., et al., A deep learning-based multi-model ensemble method for cancer prediction. Comput Methods Programs Biomed, 2018. 153: p. 1–9.

92. Cortes-Ciriano, I. and A. Bender, KekuleScope: prediction of cancer cell line sensitivity and compound potency using convolutional neural networks trained on compound images. J Cheminform, 2019. 11(1): p. 41.

93. Zhao, T., et al., Identifying drug-target interactions based on graph convolutional network and deep neural network. Brief Bioinform, 2020.

94. Huang, C., et al., Open source machine-learning algorithms for the prediction of optimal cancer drug therapies. PLoS One, 2017. 12(10): p. e0186906.

95. Ozturk, H., A. Ozgur, and E. Ozkirimli, DeepDTA: deep drug-target binding affinity prediction. Bioinformatics, 2018. 34(17): p. i821–i829.

96. Liu, Z., et al., DeepScreening: a deep learning-based screening web server for accelerating drug discovery. Database (Oxford), 2019. 2019.

